# Disrupted maternal care alters neural-microglia interactions in the primate paralaminar (PL) nucleus of the amygdala

**DOI:** 10.64898/2026.01.06.697965

**Authors:** Dennisha P. King, Mayesa Khan, Ania K. Majewska, Judy L. Cameron, Julie L. Fudge

**Author notes:** Corresponding author: Julie L. Fudge, MD Department of Neuroscience, University of Rochester Medical Center Rochester, NY 14642. Conflicts of Interest: The authors report no conflicts. Grant Support: Schmitt Program for Integrative Neuroscience at the Del Monte Institute for Brain Science (URMC), Intellectual Developmental Disability Research Center Cell and Molecular Imaging Core (URMC) (P50 HD103536), the National Institute of Neurological Disorders NS115705 (T32 NS115705 D.P.K), R21MH127486 (JLC, JLF). **Author contributions**. DK performed research, analyzed the data, and wrote the first draft of the paper. MK performed stereology research. JLC designed behavioral studies and provided editorial input, AM provided expertise on data interpretation and edited the paper, JF designed the research, supervised data analysis of anatomic data, and edited the paper.

## Abstract

Prolonged postnatal maturation of the primate amygdala is thought to be driven, at least partially, by continued neural maturation within the paralaminar nucleus (PL). At birth, the PL is densely populated with post-mitotic glutamatergic neurons that gradually mature throughout postnatal life. This active process is likely supported by microglia, which promotes synaptic maturation. Our previous work showed that early life stress associated with maternal separation alters microglia development across the infant to adolescent transition. Here, we examined whether these morphologic microglial changes are associated with alterations in the numbers of pre-synaptic terminals (SYN1+ puncta), post-synaptic terminals (PSD95+ puncta), and putative excitatory contacts (SYN1-PSD95 colocalization), and whether these synaptic elements are engulfed by phagocytic microglia. In maternally reared animals, SYN1+ puncta, PSD95+ puncta, and putative synaptic contacts decreased, while microglial (IBA1+) volume, CD68+ content, and engulfment of synaptic elements increased, between infancy and adolescence. These findings suggest greater pruning of all synaptic elements by adolescence. Maternal separation altered this trajectory, resulting in increased phagocytic activity and engulfment of synaptic elements in infancy, but not in adolescence. Maternal separation also resulted in a 50% reduction in mature PL neurons by adolescence, suggesting maturational failure, cell loss, or both by adolescence. These findings demonstrate that early life stress disrupts normative synaptic pruning and microglia-synapse interactions in the developing primate PL. Increased synaptic engulfment in infants with disrupted care is associated with premature, aberrant pruning, and highlights a potential cellular mechanism through which early environmental insults could change PL neural development by adolescence.

**Significance Statement:** The paralaminar nucleus (PL) of the amygdala is an important substrate for the delayed post-natal development of the amygdala in human and nonhuman primates. Gradually maturing glutamatergic neurons in this region, and the microglia that support them, are exposed to life events which may shape their development. We recently found that maternal separation in infants produces aberrant hyper-ramified microglia in the PL beginning in infancy and persisting into adolescence. Examining neuron-microglial interactions in the same cohort, we now find Increased phagocytic engulfment of synaptic elements by microglia after maternal separation in infancy only, with a reduction in PL mature neurons that is apparent by adolescence. Together these data suggest a mechanism for altered PL maturation, instigated by disrupted maternal care.

---

In the human and nonhuman primate, the amygdala undergoes extensive structural and functional development from infancy through adolescence supporting the emergence of affective and social behaviors (Tottenham and Sheridan, 2009; Payne et al., 2010; Goddings et al., 2014; Schumann et al., 2019). The amygdala’s paralaminar nucleus (PL) stands out as a unique substrate for amygdala growth as it contains hundreds of thousands of immature glutamatergic neurons that shift gradually toward a higher proportion of mature neurons by adolescence (de Campo et al., 2017; Avino et al., 2018; Sorrells et al., 2019; Chareyron et al., 2021; Page et al., 2022; McHale-Matthews et al., 2023).

One key process supporting the maturation of immature neurons is synaptic pruning by microglia: the selective elimination of excess or weak synapses which refines neuronal connectivity and shapes functional circuits (Rakic et al., 1986; Paolicelli et al., 2011; Petanjek et al., 2011). Microglia-mediated pruning and remodeling of synaptic elements refines dendritic architecture and strengthens functional connectivity across maturing neural networks. Thus, synapse formation and elimination are influenced not only by the timing of afferent neuronal activity but also by microglia-mediated mechanisms (Stevens et al., 2007; Schafer et al., 2012; Crapser et al., 2021; Menassa et al., 2022; Smail and Lenz, 2024).

We previously found that PL microglia morphology in typically developing macaques undergoes marked changes between infancy and adolescence, shifting from a characteristically amoeboid phenotype to ramified forms (King et al., 2025). We hypothesized that this transformation likely reflects increasing engagement in synaptic refinement, coinciding with known windows of neuronal maturation in the PL (Chareyron et al., 2012; McHale, Kelly and Fudge, 2017; Avino et al., 2018; Sorrells et al., 2019).

Environmental factors can shape the trajectory of microglial and neuronal development, modulating their interactions in brain regions undergoing extended maturation (Block et al., 2022). Microglia are highly sensitive to changes in their microenvironment, including immune challenges and psychosocial stress, which can alter their morphology, function, and relationship with surrounding neurons (Tremblay, Lowery and Majewska, 2010; Butovsky and Weiner, 2018; Savage, Carrier and Tremblay, 2019; Catale et al., 2020). Disrupting normative caregiver-to-infant interactions is a potent model of early life stress across species (Spencer-Booth and Hinde, 1971; Coe, Rosenberg and Levine, 1988; Sanchez, Ladd and Plotsky, 2001; Smail and Lenz, 2024). In the primate PL, where neuronal maturation and synaptic refinement are thought to extend well into adolescence, such environmental perturbations may have lasting consequences. Consistent with this idea, we previously found that maternal separation in macaques induced a hyper-ramified microglial phenotype that emerged during infancy and persisted in adolescence (King et al., 2025). These microglia exhibited increased process complexity and arborization, and enlarged somata, diverging from a trajectory of normal microglial maturation in control animals. The emergence of this hyper-ramified morphological phenotype in infancy, and its sustained presence in adolescence, is consistent with a primed microglial state, marked by heightened but dysregulated surveillance (Torres-Platas et al., 2014; Chastain et al., 2019). Dysregulated surveillance may in turn perturb the typical mechanisms of synaptic pruning and circuit refinement during development (Catale et al., 2020; Maras et al., 2022; Vidal-Itriago et al., 2022).

Shifts in microglia structure may signal disruptions in function, particularly in synaptic refinement during early neural development (Paolicelli et al., 2011; Schafer et al., 2012; Schafer, Lehrman and Stevens, 2013; Mallya et al., 2019). In the PL, where maturing glutamatergic neurons likely depend on precise microglia-mediated sculpting of connectivity, abnormal microglial engagement could contribute to long-term circuit function. Therefore, this study focuses on understanding excitatory synapse formation in the PL, and on evaluating whether developmental alterations in microglial morphology are associated with changes in the numbers of synaptic elements. We examine microglial engulfment of pre- and post-synaptic elements to determine whether early life stress modifies microglia synapse interactions in infancy and adolescence. By relating microglial structural dynamics to their roles in synaptic remodeling, we aim uncover substrates by which an important early environmental stressor can alter the trajectory of PL maturation.

## Methods

### Animals

A total of 23 rhesus macaques (*Macaca mulatta*) were included in this study (19 females, 4 males), all bred and raised at the University of Pittsburgh. These cohorts were previously used in our studies examining gene expression, neuronal density, and microglial morphology in the amygdala (Sabatini et al., 2007; de Campo et al., 2017; McHale-Matthews et al., 2023; King et al., 2025), and the husbandry and rearing paradigms described previously. Mothers and infants were socially housed in group-rearing pens containing 4-5 other individuals spanning a range of developmental stages from juvenility to adulthood. Infants were assigned to a maternally reared condition, or one of two maternal separation paradigms, and underwent bi-weekly behavioral assessments as part of broader studies of the impact of early life experience. Animals were euthanized at either 3 months of age (’infant’ group) or at 4-5 years of age (’adolescent’ group) (**Fig.1**). After euthanasia, brain tissue was transferred to the University of Rochester for histological and immunohistochemical analyses. All procedures were conducted in accordance with NIH guidelines and were approved by the University of Pittsburgh Institutional Animal Care and Use Committee.

**Figure 1:**
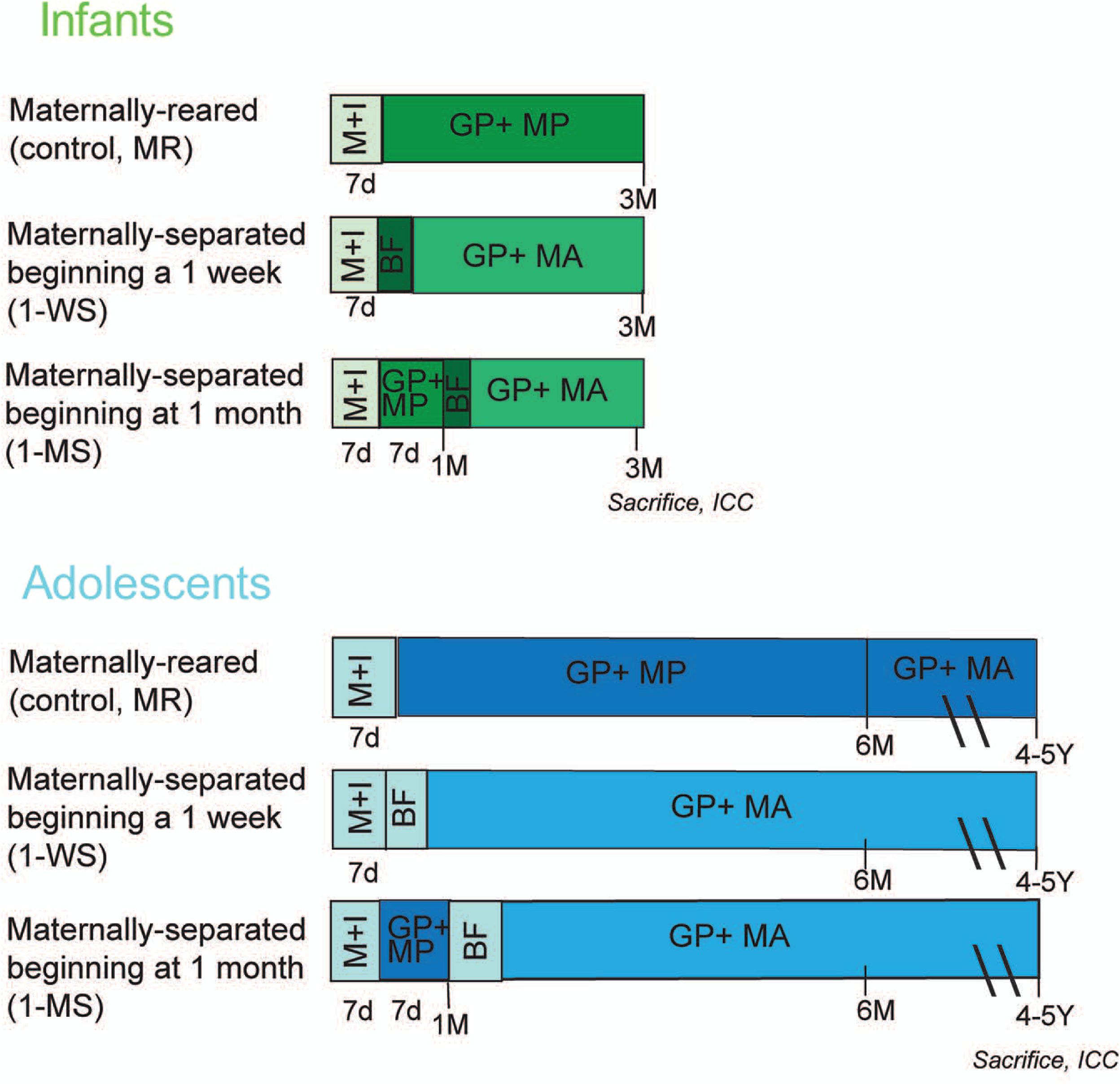
Experimental groups and housing paradigms for maternally reared (MR) controls and maternally separated (MS) animals. The experimental design spanned two developmental stages: infant cohorts (sacrificed at 12 weeks of age) and adolescent cohorts (sacrificed at 4–5 years of age). Each experimental condition was comprised of 4 animals per age group, except for the adolescent 1-WS group (n=3). Total sample size: infants (n=12), adolescents (n=11). All subjects spent the first 7 days (7d) housed with the mother-infant pair (M+I). BF, training in bottle feeding after mother removed; MA, mother absent; MP, mother present; M+I, mother-infant without group; GP, group pen environment (See text for details).

### Maternal separation paradigm

Animals were randomly assigned at birth to one of three rearing conditions: maternally reared (MR), 1-week separated (1-WS), or 1-month separated (1-MS) (**Fig.1**). 1-WS and 1-MS animals had their mothers removed at 1 week or 1 month of age, respectively, but remained in their group pens. Brain correlates of maternal separation were studied at two timepoints, resulting in an infant group (sacrificed at 3 months; n = 12, all females) and an adolescent group (sacrificed at 4–5 years; n = 11, 8 females and 3 males). All infants remained with their mothers for the first postnatal week before assignment to a rearing condition. MR animals continued to be housed with their mothers for the duration of the experiment (3 months, infant cohort) or until 6 months of age (adolescent cohort).

### Details of group-rearing environment

Macaques, like all primate species, live in interdependent social groups. Rhesus macaques have a matrilineal social structure, with females providing child-rearing (Kaufman and Rosenblum, 1967; Cameron et al., 1998; Maestripieri et al., 2006). During the first week of life each experimental infant and its mother were housed in a single cage just adjacent to a group pen. Each group-rearing pen initially housed three non-experimental female primates of juvenile to adolescent age. MR and 1-MS animals entered the group pen with their mothers in Week 2. When present, the mother assumed the dominant social role in the group, minimizing potential social stress (Sapolsky, 1996). In contrast, 1-WS animals were transitioned to bottle feeding in individual housing just outside the group pen from Days 7–14 (Similac with Iron; Abbott Laboratories) before group reintroduction at Week 2. To ease this transition, a custom-built hutch (accessible only by the separated infant) was placed in the pen, containing bottles and a soft, cloth-stuffed toy for contact comfort.

1-MS infants were similarly bottle-fed in individual cages for one week following maternal separation at Week 4 and were reintroduced to the group pen from Weeks 5-12. MR infants remained with their mothers for the full 12-week study period. Adolescents in the MR group remained with their mothers for 6 months, consistent with normative weaning in macaques, before being housed in stable, mixed-sex social groups. These adolescent groups included animals from each rearing condition to examine the social and behavioral consequences of early maternal separation. Adolescent monkeys were euthanized approximately 9 months following regrouping (mean age at sacrifice = 5.14 ± 0.17 years).

### Tissue collection and processing

Animals in each age group were euthanized under deep anesthesia and perfused with 0.9% saline. Following brain removal, brains were hemisected in the sagittal plane and blocked for flash freezing (right hemisphere) or immersion fixation (left hemisphere) (details in (Sabatini et al., 2007; de Campo et al., 2017)). Blocks of immersion fixed tissue used in this study were sectioned coronally at 40μm through the entire extent of the amygdala using a freezing sliding microtome and were stored in cryoprotectant in serial compartments.

### Confocal studies

#### Immunohistochemistry

We used triple and quadruple immunofluorescence staining in free-floating tissue in these studies. Antibodies were chosen due to their prior validation in rodent and primate species (Mouton, Price and Walker, 1997; Karube, Kubota and Kawaguchi, 2004; Innocenti and Caminiti, 2017) (**Table 1**). In the first studies, pre-synaptic elements were identified using antibodies to synapsin 1,SYN1, a presynaptic vesicle protein exclusively associated with small vesicles in neuron terminals (Navone, Greengard and De Camilli, 1984)). Excitatory post-synaptic elements were identified with anti-sera to postsynaptic density-95, PSD-95, a postsynaptic density scaffolding protein specific to excitatory synapses (Subramanian et al., 2019). Localization of these puncta and their colocalization within the PL was accomplished using doublecortin (DCX, a marker of immature neurons) (Gleeson et al., 1999)) to identify the DCX-enriched cell somas typical of the PL. A second series of adjacent sections were quadruple fluorescence immunolabeled using antibodies directed against: Ionized calcium-binding adaptor molecule 1 (IBA1) (a microglial marker (Ito et al., 1998; Ferrara et al., 2022)), the human-specific transmembrane glycoprotein, CD68 (cluster of differentiation 68 protein, CD68)(Holness et al., 1993; Chistiakov et al., 2017) and also SYN1 and PSD95 puncta, as described above. These adjacent sections were registered to the DCX-labeled sections to localize the PL region. We then identify the colocalization of synaptic elements (SYN1+ and PSD95+ puncta) and putative synaptic contacts (SYN1-PSD95 colocalized puncta) with presumptive CD68-IBA1 co-labeled structures in the PL, thus yielding a ‘phagocytic’ index across conditions. For both experiments, amygdala sections were sampled at a 1:12 interval and run in counter-balanced batches, with the investigator blind to both age and condition of the tissue. In brief, sections were rinsed 4x 15 minutes and then overnight in 0.1M phosphate buffer (PB) containing 0.3% Triton X-100 (PB-TX). The next day the sections were treated with an endogenous peroxidase inhibitor for 5 minutes at room temperature. After six 15-minute PB-TX washes, sections were blocked in 10% normal donkey serum (NDS) in PB-TX for 30 minutes. Primary antibodies at the dilutions noted in **Table 1** were pooled in 10% NDS, in which tissue was incubated at 4°C for four nights on a rocker. Following additional rinses and blocking 10% NDS, sections were incubated in the dark with pooled and filtered species-appropriate secondary antibodies for four hours (**Table 1**). Tissue then thoroughly rinsed in 0.1MPO4 and mounted onto gelatin-coated slides. After air-drying in the dark for 2–4 days, sections were cover slipped using the aqueous medium, Prolong Gold (Thermo Fisher Scientific, Waltham, MA).

**Table 1.** Primary and secondary antibodies used in confocal and light microscopy studies.

| **Antigen / Target** | **Host Species (Isotype)** | **Vendor (Location)** | **Catalog/Lot #** | **Dilution (IHC)** |
| --- | --- | --- | --- | --- |
| ***Primary Antibodies*** |  |  |  |  |
| SYN1 (Synapsin 1) | Rabbit | Cell Signaling (Danvers, MA) | D12G5/4 | 1:2000 |
| PSD95 | Mouse | Millipore Sigma (Burlington, MA) | MABN68/3863962 | 1:10,000 |
| DCX (Doublecortin) | Guinea pig | Millipore Sigma (Burlington, MA) | AB2253/3951142 | 1:500 |
| DCX | Rabbit | Abcam (Waltham, MA) | AB18723/GR3383648 | 1:15000 |
| IBA1 | Guinea pig | Synaptic Systems (Göttingen, Germany) | 234-308/1-17 | 1:5000 |
| CD68 | Goat | R&D Systems (Minneapolis, MN) | AF2040/KOR026121 | 1:2000 |
| ***Secondary Antibodies*** |  |  |  |  |
| Donkey anti-Guinea pig IgG (DyLight 405) | Donkey | The Jackson Laboratory (Bar Harbor, ME) | 706-475-148/166078 | 1:200 |
| Goat anti-Guinea pig IgG (Alexa Fluor 488) | Goat | Thermo Fisher Scientific (Waltham, MA) | A11073/2160428 | 1:200 |
| Donkey anti-Rabbit IgG (Alexa Fluor 488) | Donkey | Thermo Fisher Scientific (Waltham, MA) | A21206/2873188 | 1:200 |
| Donkey anti-Goat IgG (Alexa Fluor 546) | Donkey | Thermo Fisher Scientific (Waltham, MA) | A11056/1216117 | 1:200 |
| Donkey anti-Mouse IgG (Alexa Fluor 647) | Donkey | Thermo Fisher Scientific (Waltham, MA) | A31571/2892424 | 1:200 |

#### Image acquisition

Images from immunofluorescent labeled slides were collected using the Nikon A1R HD Laser Scanning Confocal with NIS-Elements software (Center for Advanced Light Microscopy and Nanoscopy). Overview images using a 4x/0.10 NA Nikon Plan Apochromat VC objective were first completed to locate regions of interest (ROI’s) to identify the boundaries of the PL. We used either the DCX-labeled channel alone, or for quadruple labeled adjacent sections, landmarks associated with DCX-labeled PL from adjacent sections. Three ROIs were aligned over the PL, and marked (medial, central, and lateral) under 2x, and high-resolution (60x) Z-stack images were then collected using the 60x oil objective (**Fig. 2A**). High power image stacks were collected at 0.125μm intervals through a Z plane (10μm plane) at a resolution of 2048 × 2048 pixels. A total of 9 ROIs were collected for each experimental animal (rostral, central, and caudal sections x medial, central, and lateral R0Is). Excitation and emission settings were as follows: DyLight 405, ex 405 nm, em 420-480 nm; Alexa Fluor 488, ex 488 nm, em 525/50 nm; Alexa Fluor 546, ex 561 nm, em 595/50 nm; and Alexa Fluor 647, ex 640 nm, em 650–700 nm. Detector gain, pinhole size, and laser power were kept constant across all samples within an experiment.

**Figure 2:**
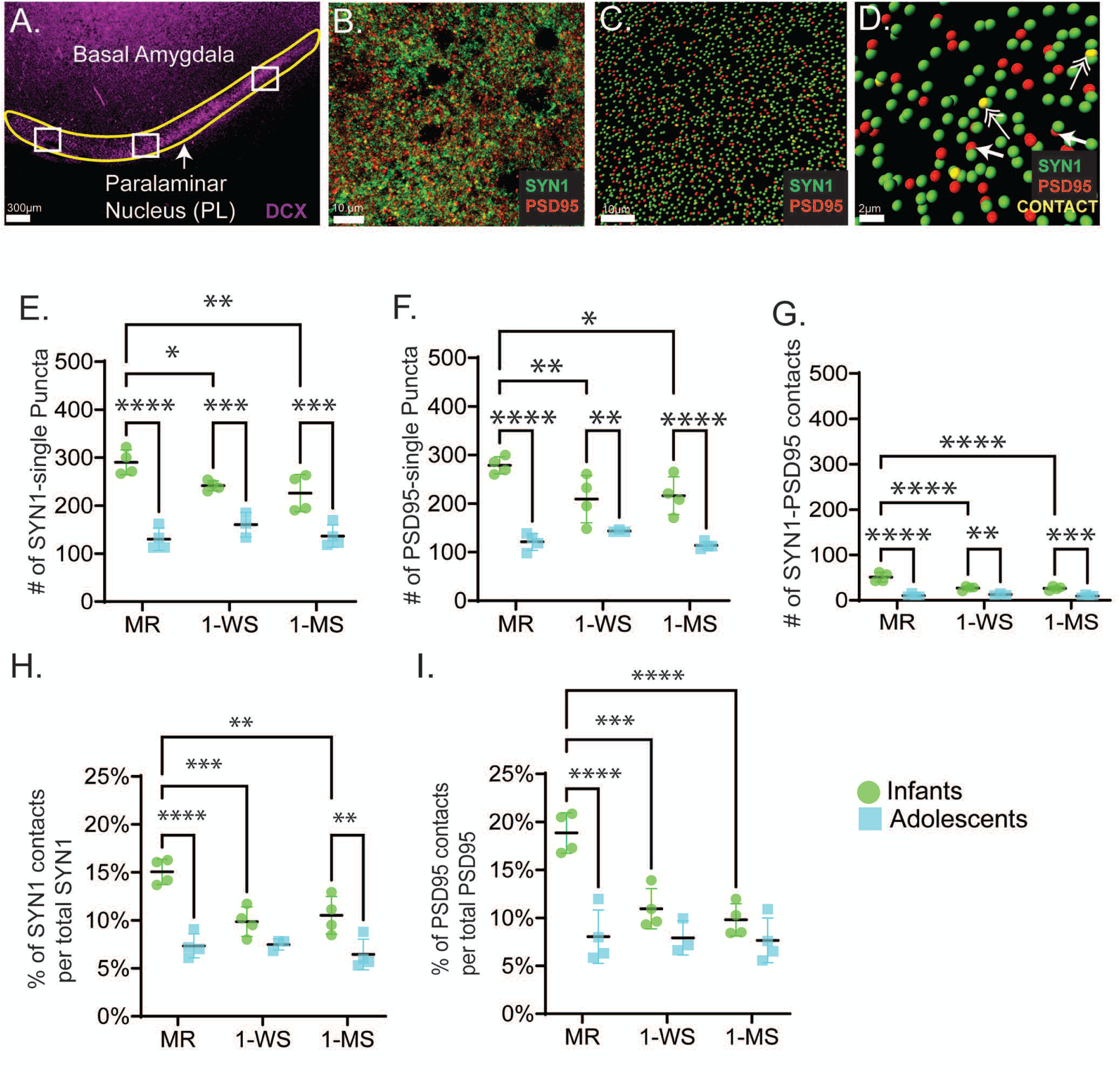
Developmental and maternal separation effects on synaptic puncta in the PL. **A.** Low-magnification (4x) overview of the paralaminar nucleus (PL) ventral to the basal amygdala, highlighting the location of the analysis identified by (DCX+) neurons. Scale bar: 300 µm. **B.** High-magnification (60x) photomicrograph of the PL neuropil depicting SYN1(green) and PSD95 (red) immunoreactivity. Scale bar: 10 µm. **C.** Spot rendering of SYN1 (green) and PSD95 (red) puncta at low power. Scale bar: 10 μm. **D.** Putative synaptic contacts (SYN1 and PSD95 overlap, greater than 50%) in yellow, double arrow. Single arrows show SYN1 and PSD95 puncta pairs that are non-overlapping. Scale bar = 2 μm. **E.** Mean number of SYN1-single labeled puncta across groups. **F.** mean number of PSD95-single labeled puncta across groups. **G.** Mean number of putative synaptic contacts (SYN1-PSD95 colocalized) across groups. **H.** Percentage of SYN1 contacts per total available SYN1+ puncta. I. Percentage of PSD95 contacts per total available PSD95+ puncta. Statistical significance: *p < 0.05, **p < 0.01, ***p < 0.001, ****p < 0.0001.

#### Analyses

Image stacks for all experiments were analyzed with Imaris 10.1 software (Bitplane). To ensure there would be no penetration issue biasing the results, all stacks were trimmed to a final height of 3μm (24 steps total) after centering the stack in the region of optimal immunostaining for all antibodies. Image stacks were then cropped in the X (150μm) x Y (150μm).

#### SYN1-PSD95 spot analysis

To analyze the synaptic elements of the PL, we used the Imaris “spot rendering” module, with sensitivity thresholds determined from interactive histograms to capture as many discrete puncta as possible (**Fig. 2B-C**). SYN1+ presynaptic puncta have been reported to vary widely in size depending on species, brain region, and method of measurement. Across cortical and hippocampal regions, presynaptic puncta typically range from ∼0.2 to 1.0µm³ in volume (∼0.5 - 1.5µm in diameter when approximated as spheres), with larger puncta occasionally reaching several µm³ in pyramidal neurons (Murthy, Sejnowski and Stevens, 1997; Schikorski and Stevens, 1997; Shepherd and Harris, 1998; Rollenhagen and Lubke, 2010). On the post-synaptic side, PSD95+ labeled elements measured in fixed tissue are variable in nonhuman primate, ranging from 400-800 nm in diameter (0.4 – 0.8 µm) (Dumitriu et al., 2012). In the Imaris slice mode, we employed the line measurement tool to randomly assess the cross-sectional diameters of SYN1+ puncta and PSD95+ puncta. Based on these measurements, we applied a 1 µm maximum XY spot diameter for SYN1 and PSD95 puncta in Imaris. This parameter aligns with the confocally resolvable dimensions of puncta and PSD clusters and retains the ability to detect smaller structures whose intensity distributions are captured within a 1 µm kernel. Puncta larger than 1 µm in diameter were excluded from the analysis. This approach balances biological accuracy with the resolution constraints of confocal imaging and is consistent with previous studies applying spherical spot models to synaptic puncta segmentation. Results for ‘spot rendering’ are reported as number of spots within the ROI. Mean volumes for ‘spots’ showed no significant differences among groups and were not further analyzed (SYN1, infants = 1035 ± 114.2µm^3^, adolescents = 1079 ± 140.0 µm3; p=0.6057; PSD95, infants = 974.4 ± 122.3µm^3^, adolescents = 988.9 ± 207.8µm^3^; p=0.8744).

We implemented a confocal microscopy strategy to evaluate the proximity and organization of structures considered putative contacts (**Fig. 2D**). While electron microscopy is necessary to confirm true synaptic contacts, estimating putative contacts can be done using several techniques. False-positive contacts can be controlled by requiring a minimum number of overlapping voxels for an object to be classified as a true contact (Wouterlood et al., 2007). We used the criteria for putative contacts based on a greater than 50% colocalization of SYN1 and PSD95 rendered ‘spots’ (**Fig. 2D**, double arrows, yellow). “Contacts” thus were determined by measuring the distance from the center of a spot (1µm diameter maximum) to the outside boundary of another rendered spot. Our analysis was restricted to a maximum object to surface distance of 0.5μm (at least a 50/50 overlap of spot and surface objects) to create a relatively stringent inclusion criteria for assessing “synaptic contacts.”

#### Engulfment studies (IBA1-CD68-SYN1-PSD95)

We employed two complementary analytical approaches to capture microglia-synapse interactions, balancing cell-level specificity with neuropil-level sensitivity. The first approach involved rendering 10 individual microglia per animal to quantify mean IBA1 volume, CD68 colocalization within IBA1-labeled microglia, and the engulfed synaptic elements on a per-cell basis. This high-specificity method allowed us to capture the full morphological diversity of microglia; including the soma and attached processes, and to directly assess phagocytic activity in fully reconstructed cells. Each microglia selected had a visible cell body attached to extensive processes and was rendered with a smoothing of 0.275μm and a local background subtraction filter with a radius of 15μm, and threshold values were applied uniformly across all image stacks. CD68+ structures associated with the rendered IBA1+ cells were rendered with a smoothing of 0.01μm and the same background subtraction, with thresholds uniformly applied. CD68 is a heavily glycosylated lysosomal/endosomal transmembrane protein that has a long, folded extracellular membrane exposed tail (Holness et al., 1993). This extracellular tail is critical for antigen detection and is frequently apparent in 60x images (**Fig. 3A, 3E, insets).** We measured both the CD68 + elements colocalized within the IBA1 volume and the contiguous CD68 segment that traversed the membrane into the extracellular space, when present. All CD68+ elements are referred to microglia (IBA1)-associated CD68 elements and had either partial or complete co-localization with the IBA-1 volumes. SYN1+, PSD95+ and putative ‘contacts’ were then characterized as ‘engulfed’ if they colocalized with any portion of IBA-1-associated CD68 elements.

**Figure 3:**
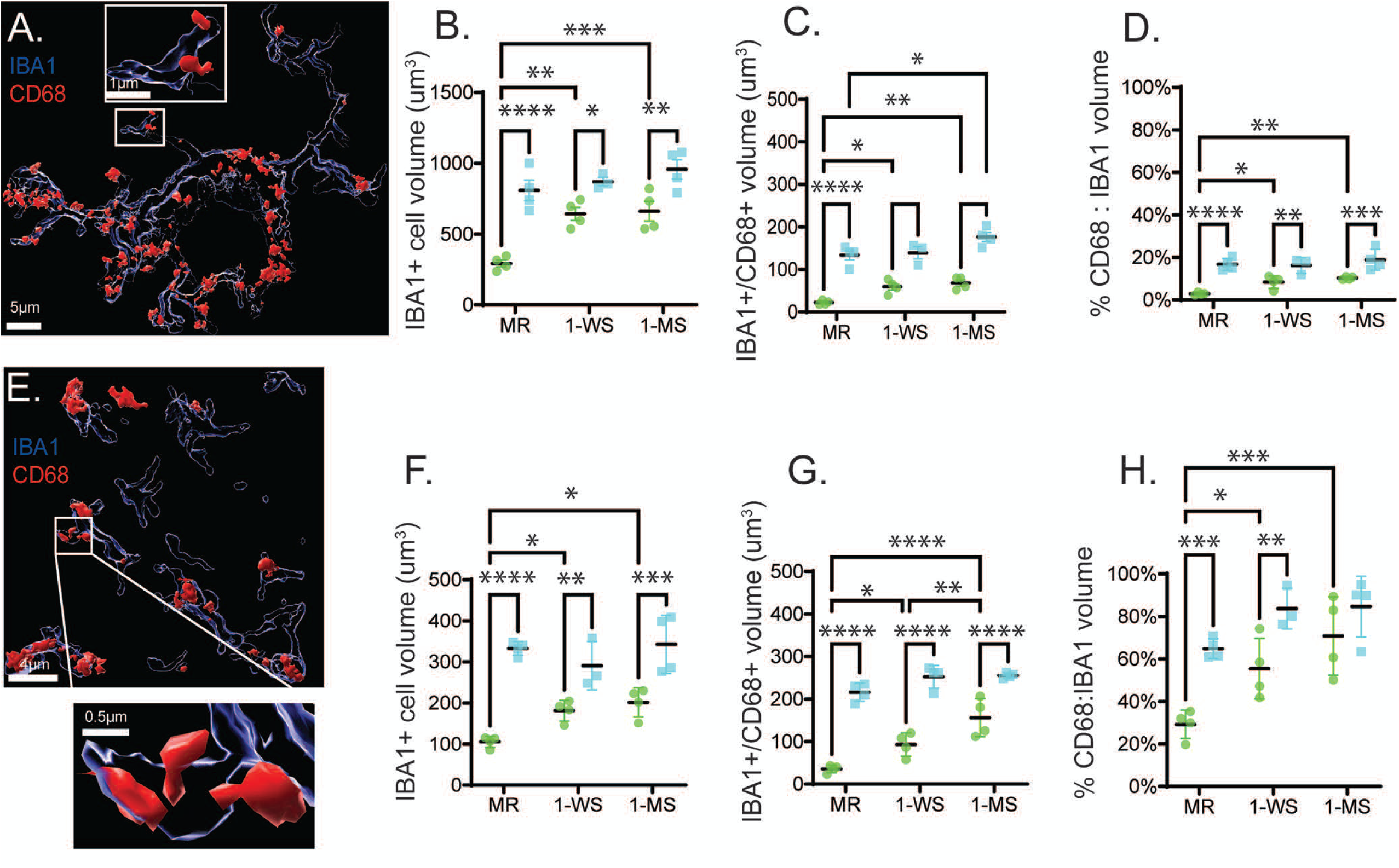
Microglial morphology (IBA1 volume) and phagocytic capacity (CD68 volume and density) in the PL. **A.** Representative image illustrating reconstruction of individual microglia, IBA1+ microglia (blue) and CD68 (red) that overlaps IBA1 volumes (see methods); inset shows examples of CD68 transmembrane colocalization in processes. **B-D**. Quantitative results from the microglia isolation method. **B.** Mean volume of IBA1+ cells (µm^3^). **C.** Mean volume of microglial-associated CD68 per IBA1 cell (µm^3^). **D.** Ratio of the microglia-associated CD68 volume to IBA1 cell volume, expressed as percent. **E-F.** Quantitative results from the ROI neuropil analysis. **E.** IBA1+ in ROI (blue) and CD68 (red) colocalization. Inset shows examples of CD68 transmembrane colocalization. **F.** Mean IBA1 volume per ROI. **G.** Mean colocalized CD68 per ROI. H. Ratio of the volume of CD68 colocalized IBA1 to total IBA1 volume per ROI expressed as a percentage. Statistical significance: *p < 0.05, **p < 0.01, ***p < 0.001, ****p < 0.0001.

In the second approach, we used a region-of-interest (ROI) method to generate a different phagocytic index within a volume of 3×10⁵ µm³ of PL neuropil, captured at 60x. Here we excluded microglial cell soma, and focused only on densely sampled neuropil, hypothesizing that this approach might be more sensitive to the ‘early’ stages of the phagocytic activity where synaptic elements are first detected and engulfed (Sierra et al., 2013; Villani et al., 2019). This ROI approach also permitted a comparison of the proportion of ‘engulfed’ versus ‘un-engulfed’ synaptic elements in the neuropil in an unbiased manner. 10 randomly selected ROI volumes per animal were captured. Using surface rendering, we quantified the number of SYN1+, PSD-95+, and putative synaptic contacts engulfed by microglia (i.e., colocalized with IBA1-associated CD68+ elements) in each ROI per animal. SYN1+, PSD95+ and putative ‘contacts’ within the microglia were then characterized as ‘engulfed’ or not.

### Stereology of immature and mature neuron counts

#### Immunocytochemical preparation for stereology

We analyzed changes in immature (DCX+) and mature (Nissl-stained pyramidal neurons) across all animals. Our results in the infant cohorts were previously published (McHale-Matthews et al., 2023); here we assessed the adolescent cohorts. In brief, compartments (1:12 series) for each animal contained evenly spaced sections with a random starting point. All staining batches were counterbalanced in a blind manner. *Doublecortin (DCX).* Conditions for DCX immunostaining were first established in the control animals that were not part of this study (Fudge, deCampo and Becoats, 2012). Tissue was rinsed in PB with 0.3% Triton-X (PB-TX) overnight. The next day, brain slices were treated with an endogenous peroxidase inhibitor for 5 minutes and then underwent more rinses in PB-TX. Sections were then pre-incubated for 30 minutes in 10% normal goat serum blocking solution with PB-TX (NGS-PB-TX). All sections were then incubated in primary antisera to DCX (1:15000, Abcam, rabbit), at 4°C for four nights. Sections were then thoroughly rinsed, blocked with 10% NGS-PB-TX, and incubated for 40 minutes in the appropriate biotinylated secondary antibody. After more rinses, sections with bound anti-DCX antibodies were incubated in an avidin-biotin complex (Vectastain ABC Elite; Vector Laboratories), visualized with 3,3’-Diaminobenzidine (DAB), and then activated with 0.3% hydrogen peroxide (H_2_O_2_). DCX-stained sections were mounted onto gelatin coated slides from 0.1M PB solution and air-dried over a 2-to 4-week period. They were then rapidly dehydrated and rehydrated, and counterstained with a light cresyl violet stain (Chroma-Gesellschaft; West Germany), and cover slipped with DPX Mounting Medium (Electron Microscopy Sciences; Hatfield, PA). To maintain the section height required for optical fractionator analyses, slight modifications to the Nissl-staining protocol were made to minimize dehydration from ethanol.

#### Optical fractionator analyses

Neuron counts were estimated using an unbiased, systematic, random sampling method known as the optical fractionator (Gundersen and Jensen, 1987; West, Slomianka and Gundersen, 1991) (Stereoinvestigator, Microbrightfield Biosciences, Williston, VT). Briefly, the PL was outlined under low power magnification (2x objective) (McHale-Matthews et al., 2023). Sampling parameters, such as counting frame and scanning grid dimensions, were first established for each neural population by oversampling to establish a coefficient of error (CE) <0.10. All DCX-positive neurons were counted as “immature” neurons, regardless of morphology; “mature” neurons were defined as DCX-negative cresyl violet stained cells with characteristic nuclear staining (Chareyron et al., 2012; Garcia-Cabezas et al., 2016). Section thickness was collected at every sampling site, so that the final cell estimates were calculated using “number weighted section thickness”. 1:12 sections double labeled for DCX and Nissl were examined with an Olympus UPlanFL 100x/1.30 oil lens using an Olympus AX70 microscope interfaced with Stereoinvestigator via a video CCD (Microbrightfield, Williston, VT). On mean, 8 sections per animal were examined (range = 7-9 sections) with approximately 480mmly of distance between each slide.

### Statistics

All statistical analyses were carried out in GraphPad Prism (version 10.5.0; GraphPad Software, San Diego, CA) for Windows. A two-way ANOVA was used to compare the developmental and experimental condition differences in the total number of SYN1, PSD95, and colocalized puncta, volume of microglia, volume of CD68 within IBA1, and volume of engulfed synaptic elements and putative synaptic contacts. Multiple comparisons were adjusted using Tukey’s multiple comparison test and are given in the text. Cell counts of immature and mature neurons were assessed using a one-way ANOVA to test the effects of maternal separation in infancy and adolescence. Statistical significance was set at p < 0.05, and all error bars represent the standard deviation (SD).

## Results

### Synaptic profiles in infants and adolescents

To provide an overview of synaptic profiles during normal PL development, we first quantified the number of presynaptic elements between maternally reared (MR) infants and adolescent macaques per ROI (3x10^4^µm^3^) (**Fig. 2A-G**). ‘Pre-synaptic’ and ‘post-synaptic’ elements were defined as elements not engaged in putative contacts. The total number of pre-synaptic elements (SYN1+ single labeled puncta), excitatory post-synaptic elements (PSD95+ single labeled puncta) and putative synaptic contacts (SYN1+-PSD95+ contacts) per ROI in the PL were all significantly decreased in the MR adolescents (mean= 130.1 ± 23.39; 121.1 ± 17.46; 10.49 ± 3.759, respectively) compared with MR infants (mean = 290.1 26.15; p<0.0001, 278.5 ± 17.63; p < 0.0001, 51.09 ± 10.57; p < 0.000, respectively)(**Fig. 2E-G**).

To determine whether the number of putative excitatory contacts was primarily constrained by presynaptic or postsynaptic availability, we normalized putative SYN1-PSD95 colocalized contacts to either presynaptic (total contacts/SYN1) or postsynaptic (total contacts/PSD95) abundance. We found that percentage of contacts per putative presynaptic (SYN1) and postsynaptic (PSD95) elements were both markedly higher in infants compared with adolescents (**Fig. 2H-I**). Control infants showed more than a two-fold greater number of both SYN1 (15.05 ± 1.311% versus 7.332 ± 1.246%; p<0.0001) (**Fig. 2H**) and PSD95 puncta (18.85 ± 2.114% versus 8.030 ± 2.786% p<0.0001) (**Fig. 2I**) engaging in contacts relative to adolescents.

### Maternal separation disrupts the synaptic profile of the PL during infancy

On the pre-synaptic side, maternal separation resulted in markedly reduced SYN1+ only puncta in both 1-week separation (1-WS) (241.5 ± 10.43; p= 0.0429) and 1-month separation (1-MS) (225.9 ± 38.79; p=0.0075) infant animals compared with maternally reared (MR) infant controls (290.1 ± 26.15; **Fig. 2E**). In contrast, no significant differences were observed in the number of SYN1+ presynaptic elements across adolescent groups (MR adolescent group [130.1± 23.39], 1-WS adolescents [160.5 ± 25.82, p= 0.3024]; 1-MS adolescents [136.5 ± 23.23, p= 0.9358]; **Fig. 2E**). PSD95+ only elements also declined in infants in both the 1-WS (209.1 ±48.81, p=0.0081) and 1-MS (216.2 ± 38.92, p=0.0169) groups in comparison with the MR group (278.5 ± 17.63, **Fig. 2F**). Amongst the adolescent groups, no significant differences were observed in the number of PSD95+ postsynaptic elements. The MR adolescent group had 121.1 ± 17.46 PSD95+ puncta, compared with 1-WS adolescents (143.4 ± 2.385, p=0.5715) and 1-MS adolescents (113.8± 7.335, p= 0.9306; **Fig. 2F**).

In infants, maternal separation significantly reduced the number of putative synaptic contacts (SYN1-PSD95 overlap), with 1-WS animals having a mean of 26.95 ± 4.903 (p<0.0001) and 1-MS animals 26.29 ± 4.961 (p<0.0001) in the PL, compared with 51.09 ±10.57 in MR controls (**Fig. 2G**). As with pre- and post-synaptic puncta, there were no observed differences between the number of putative synaptic contacts between the 1-WS (12.97 ± 2.386, p=0.8398) and 1-MS adolescent groups (9.431 ± 2.863 p= 0.9634) and MR adolescent group (10.49 ± 3.759; **Fig. 2G**).

The percentage of contacts per total available SYN1 elements declined in infants (1-WS, 9.859 ± 1.544%, p=0.0003, 1-MS, 10.52 ± 1.956%, p= 0.0012) compared to MR controls (15.05 ± 1.311%). This ratio was not affected in the adolescent groups (MR (7.332 ± 1.246%),1-WS (7.478 ± 0.5734%, p=0.9907), and 1-MS (6.443 ± 1.598%, p=0.6751) (**Fig. 2H**). For PSD95, the percentage putative contacts shifted in maternally separated infants, decreasing the proportion of contacts made per the available PSD95 pool (MR, 18.85 ± 2.114%, 1-WS, 10.94± 2.101, p=0.0002, 1-MS, 9.792± 1.687, p<0.0001). This did not change in the adolescent groups (MR, 8.030 ± 2.786, 1-WS, 7.917 ± 1.782, p=0.9975, 1-MS, 7.647 ± 2.330, p=0.9668; **Fig. 2I**). Together, these results suggest there were fewer synaptic contacts per available pool of presynaptic and post-synaptic elements infants, bringing the rate closer to adolescent levels.

### Microglia volume increases between infancy and adolescent in normal PL

There was a 176.023% increase in the mean volume of microglia between infancy (293.2 ± 47.97µm^3^) and adolescence (809.3 ± 143.5 µm^3^, p<0.0001; **Fig. 3A-B**, mean volume of IBA1 labeled cells, n=10 per animal) in MR groups, in general agreement with our previous morphologic results using different methods (King et al., 2025). Increased CD68 is often used as a measure of presumed ‘phagocytic’ activity in post-mortem brain (Chistiakov et al., 2017). Although this measure requires ‘real-time’ visualization of phagocytosis, we used ex vivo measures to approximate net phagocytic activity in the PL across conditions (**Fig. 3C**). The mean volume of microglia-associated CD68 between control infants and adolescents showed a substantial increase (507.8% change) (MR infant, 21.98 ± 4.795 µm^3^) and adolescent (MR adolescent 133.6 ± 22.60 µm^3^, p< 0.0001; **Fig. 3C**). Normalizing CD68 volume to IBA1 cell volume per cell, yielding a measure of lysosomal ‘density’ per microglia. The MR adolescent group (16.68 ± 2.845%) exhibited a significantly higher CD68/IBA1 ratio compared to the MR infant group (2.850 ± 0.5859%, p<0.0001; **Fig. 3D**), indicating a developmental increase in CD68 driven by relative concentration per microglial cell, beyond an increase in overall microglial size.

### Maternal separation increases the volume of microglia and phagocytic activity during infancy but not adolescence

IBA1 volume was increased in both maternally separated infant groups (1-WS, 642.6 ± 93.08 µm^3^, p=0.0010; 1-MS, 661.4 ± 138.0 µm^3^, p=0.0006) relative to MR infant animals (293.2 ± 47.97µm^3^, **Fig. 3B, green**). However, there were no significant differences in the IBA1 volume between the 1-WS (869.9 ± 50.48 µm^3^, p= 0.7613) and 1-MS (957.2 ± 136.3 µm^3^, p=0.1777) adolescent animals in comparison to the MR adolescent control group (809.3 ± 143.5 µm^3^; **Fig. 3B, blue**).

In the infant group, we observed robust increases in the volume of CD68 within IBA1+ cells, in both maternally separated groups (1-WS (59.24 ± 15.93 µm^3^, p=0.0260) and 1-MS (67.79 ± 14.44 µm^3^, p=0.0065) relative to the MR group (21.98 ± 4.795 µm^3^; **Fig. 3C**). Whereas there were no observed differences between the MR adolescent group (133.6 ± 22.60 µm^3^) and the 1-WS adolescent group (138.8 ± 25.04 µm^3^, p= 0.9288), there was a significant increase between the MR and 1-MS adolescent group (176.3 ± 21.60 µm^3^, p=0.0110; **Fig. 3C**). The elevated CD68 levels in both maternally separated groups disrupted the significant increase normally observed between infancy and adolescence.

Taking the percentage of IBA1 volume occupied by CD68 for every cell rendered, we found that the ‘density’ of CD68 per microglia was also affected by maternal separation in the infants (1-WS, 8.350 ± 2.760%, p=0.0473,1-MS,10.25 ± 0.6019%, p=0.0075), but not adolescent groups (1-WS, 16.20 ± 3.751%, p= 0.9766, 1-MS,18.90 ± 4.882%, p= 0.5581; **Fig. 3D**). Thus, the rise in CD68 with age in MR animals (**Fig. 3B**), may reflect maturational changes in microglial morphology (**Fig. 3A**) as well as upregulation of phagocytic capacity. The same may also be true in infants affected by maternal separation.

#### ROI approach

Using our complementary method measuring IBA1 and CD68 volumes within a neuropil ROI (3 x 10^4^ mm^3^), we found generally similar results (**Fig. 3E-G**). While this approach did not provide information on individual microglial morphology, it provided a view of ‘process-oriented’ phagocytosis, an early phagocytic event (Sierra et al., 2013; Stotzel and Kiermaier, 2022). Compared to results of the individualized microglial rendering study (**Fig. 3A-C**), there were similar increases in the mean volume of IBA1 across MR control groups (infants: 105.5 ± 13.70µm^3^ versus adolescents: 332.8 ±16.74µm^3^, p <0.0001, **Fig. 3F**), and also significant increases in microglial volume in both infant maternally separated groups in comparison with the MR control (1-WS, 181.4 ± 25.28 µm^3^, p= 0.0471,1-MS, 201.5 ± 35.58 µm^3^, p= 0.0116). Consistent with the individualized microglia approach (**Fig. 3B**), there were also no changes in microglial volume amongst the adolescent maternally separated groups (1-WS, 290.9 ± 58.91 µm^3^, p= 0.3995,1-MS, 342.7 ± 70.68 µm^3^, p= 0.9391) compared to the MR adolescent control group (**Fig. 3F**).

Microglia-associated CD68 volumes (**Fig. 3G**) were also significantly increased in MR infant (35.52 ± 8.987µm^3^) versus MR adolescent PL (216.0 ± 21.15µm^3^, p <0.0001). Maternal separation greatly increased the volume of colocalized CD68 with IBA1 in the neuropil in both 1-WS infants (92.92 ± 27.46µm^3^, p= 0.0154) and 1-MS infants (156.1 ± 44.88µm^3^, p <0.0001). Adolescent CD68 volumes were not significantly altered by maternal separation conditions (1-WS, 252.6 ± 26.77µm^3^, p= 0.1435, 1-MS, 255.6 ± 6.503µm^3^, p= 0.1072; **Fig. 3G**), reproducing the general results in the whole microglia approach.

‘Density’ ratios for CD68-IBA1 colocalized volumes in the ROI neuropil method largely replicated those in the individual microglial analysis (**Fig. 3D** versus **Fig. 3H**), with an increased in the ratio in MR infants (29.21 ± 6.687%) compared to MR adolescents (64.85 ± 4.607%, p= 0.0008). Similarly, maternally separated infants, both 1-WS (55.43 ± 14.29%, p=0.0218) and 1-MS (70.81 ± 18.45%, p=0.0005) had higher CD68:IBA1 volume ratios in comparison to the MR infant control (29.21 ± 6.687%). Again, there were no significant differences amongst the maternally separated adolescent groups versus adolescent MR control (1-WS, 83.67 ± 9.629%, p=0.1476, 1-MS 84.64 ± 14.33%, p= 0.0913). However, there was a loss of maturational increase in the 1-MS group (1-MS infant (70.81 ± 18.45%) compared to 1-MS adolescent group (84.64 ± 14.33%, p=0.1345), due to increased CD68 content in the infant cohort (**Fig.3H**). This pattern suggests that early life stress accelerates microglial phagocytic activity in infancy, effectively reducing the relative typical developmental change observed between infancy and adolescence.

### Increased engulfment of synaptic elements in normal development

To determine whether the observed increases in phagocytic activity corresponded to synaptic engulfment, we quantified the number of presynaptic (SYN1+) and excitatory post-synaptic (PSD95+) elements and putative synaptic contacts (SYN1-PSD95 overlapped) within microglia-associated CD68 elements in the control groups, first using the individualized microglia method (**Fig. 4A-C, D-F**). As before, ‘pre-synaptic’ and ‘post-synaptic’ elements were defined as elements not engaged in putative contacts. The number of SYN1+ elements co-localized with microglia-associated CD68 volumes (**Fig. 4A, D**) increased significantly between MR infant animals (28.025 ± 10.958) and MR adolescent animals (193.667 ± 68.742, p <0.0001). This suggested a large increase in engulfment of SYN1+ puncta by adolescence (591.051%). The number of post-synaptic PSD95+ puncta colocalized with CD68 (**Fig. 4B, E**) also dramatically increased between the MR infant (23.575 ± 9.263) and MR adolescent (177.075 ± 71.578, p<0.0001). Finally, putative synaptic contact numbers colocalized within microglia-CD68 positive structures also revealed significant increases between the maternally reared infants (9.175 ± 4.855) relative to maternally reared adolescents (91.53 ± 41.90, p<0.0001; **Fig. 4C, F**).

**Figure 4:**
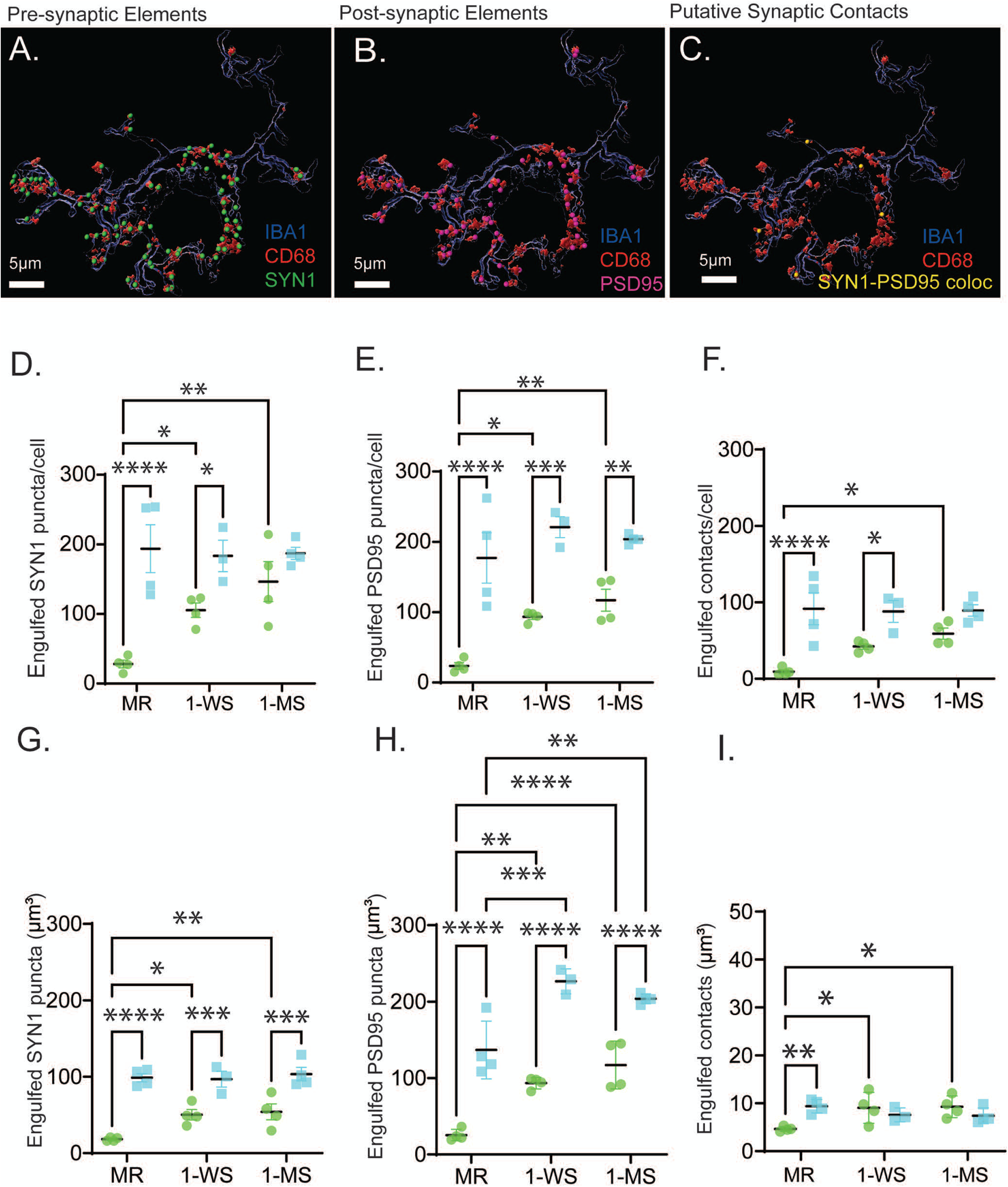
Engulfment of synaptic elements and putative contacts. **A-C**. Representative 3D spot-surface renderings for pre-synaptic elements (green)(**A.)** post-synaptic excitatory elements (pink**) (B),** and putative contacts (SYN1-PSD95 colocalized, yellow) **(C). D-F.** Individual microglia isolation method, 10 microglia/animal. **D.** Mean number of engulfed presynaptic (SYN1+) elements per microglia. **E.** Mean number of engulfed postsynaptic (PSD95+) elements per microglia. **F.** Mean number of engulfed putative synaptic contacts (SYN1-PSD95 overlap) per microglia. **G-I.** Quantitative results from the ROI neuropil analyses, 10 (3x10^4^µm^3^) ROIs/animal. **G.** Mean number of engulfed presynaptic (SYN1+) elements per neuropil ROI **H.** Mean number of engulfed postsynaptic (PSD95+) elements per neuropil ROI. **I.** Mean number of engulfed putative synaptic contacts (SYN1-PSD95 overlap) per neuropil ROI. Statistical significance: *p < 0.05, **p < 0.01, ***p < 0.001, ****p < 0.0001.

Using our ROI approach to examine the neuropil (**Fig. 4G-I**), we also found that the number of engulfed pre-synaptic elements (SYN1+) increased between the MR infant (18.24 ± 2.344 per 3x10^4^µm^3^) and adolescent groups, similar to the individual microglia method (441.118%) (98.70 ± 10.74 per 3x10^4^µm^3^, p<0.0001; (**Fig. 4G**). Analysis of post-synaptic (PSD95) elements revealed similar patterns between MR infant (25.39 ± 7.513) and MR adolescent groups (136.8 ± 37.82, p< 0.0001), with PSD95-alone engulfment increasing by 438.795% (**Fig**. **4H**). Lastly, the number of engulfed putative synaptic contacts also significantly increased (infants, 4.638 ± 0.4901 versus adolescents, 9.375 ± 1.411, p=0.0033, **Fig.4I**).

### Maternal separation increases numbers of all engulfed synaptic elements in infancy only

In the individual microglia approach, both the 1-WS (105.33 ± 20.95, p= 0.0453) and the 1-MS infant group exhibited a significant increase in the mean number of engulfed pre-synaptic elements (146.48 ± 57.41, p= 0.0025) in comparison to the MR infant group (28.025 ± 10.96; **Fig. 4D**). In adolescent groups, the mean number of engulfed pre-synaptic elements did not differ significantly among MR (193.67 ± 68.74), 1-WS (183.367 ± 38.9755, p= 0.9444), and 1-MS subjects (187.112 ± 17.436, p= 0.9733; **Fig. 4D**). Likewise, the number of engulfed post-synaptic excitatory elements (PSD95+) was also greater in both the 1-WS (93.4685 ± 7.5187, p=0.0272) and 1-MS infant groups (117.075 ± 31.1385, p= 0.0036) compared to the MR control group (23.575 ± 9.2626). In adolescent groups, the mean number of engulfed post-synaptic elements (PSD95+ puncta) was similar between the MR adolescent group (177.075 ± 71.5780) and both the 1-WS (221 ± 25.7006, p=0.2449) and 1-MS adolescent groups (203.8 ± 6.5018, p= 0.5283) (**Fig 4E**). ‘Engulfed’ putative synaptic contacts (SYN1-PSD95) were significantly elevated only the 1-MS separated group (59.03 ± 14.55, p= 0.1099) but not the 1-WS group (42.14 ± 6.865, p=0.0125) compared to the maternally reared controls (9.175 ± 4.855). Again, there were no changes in the number of engulfed contacts among the maternally separated adolescent groups (1-WS, 88.00 ± 24.75, p= 0.9754, 1-MS, 89.30 ± 14.96, p=0.09885) compared to the adolescent MR control group (91.53 ± 41.9; **Fig**.**4F**).

Our ROI approach produced many of the same findings. However, the neuropil-only analysis showed that maternal separation led to an increase in the number of engulfed pre-synaptic (SYN1+) elements in both the 1-WS infant (50.33 ± 13.47, p= 0.0191) and 1-MS infant (54.06 ± 20.82, p= 0.0091) groups in comparison to the MR infant control group (18.24 ± 2.344). There were no statistical differences among all adolescent conditions (MR, 98.70 ± 10.74, 1-WS 96.89 ± 17.91, p=0.9862, 1-MS, 103.4 ± 17.65, p=0.8957; **Fig.4G**).

In contrast, engulfed PDS95 elements in the neuropil were increased by maternal separation in both infant and adolescent groups compared to age-matched controls (infants: MR, 25.39 ± 7.513, 1-WS, 93.47 ± 7.519, p=0.0011, 1-MS, 117.1 ± 31.14, p<0.0001, adolescents: MR, 136.8 ± 37.82, 1-WS, 226.7 ±16.29, p=0.0001,1-MS, 203.8 ± 6.502, p=0.0013; **Fig.4H**). This suggests that trends observed in our individual microglial analysis were boosted to significance by biasing the analysis toward engulfment in the processes (early-stage phagocytosis). Importantly, this approach revealed increased engulfment of PSD-95 excitatory elements by microglial processes in adolescents that experienced maternal separation years prior.

Lastly, evaluation of the neuropil revealed that engulfed putative synaptic contacts in MR adolescents were higher than in MR infants, while maternal separation increased engulfment only in infant groups (infant MR,4.638 ± 0.4901 versus 1-WS, 9.044 ± 3.222, p=0.0145 and 1-MS, 9.269 ± 2.250, p=0.0103; adolescent MR control, 9.375 ± 1.411 1-WS 7.575 ± 1.453, p=0.4684, 1-MS, 7.375 ± 1.650, p=0.3427) (**Fig. 4I**). These results are generally similar to the whole microglia assessment. However, we found increased engulfment of putative contacts in both infant separated groups compared to the MR infant, a result not seen in the individualized microglial assessments (**Fig. 4F**).

### Maternal separation results in decreased mature neurons by adolescence

In our previous investigation quantifying neuron changes in the infant cohort only, there were no changes in immature or neuron numbers (McHale-Matthews et al., 2023) (**Fig. 5A, B**), despite the subtle changes in immature neuron soma volume in the infant separation groups, an indicator of neural maturation (Ryan, Ehrlich and Rainnie, 2014). In that study, we hypothesized that there were few changes in infants, which were sacrificed 4-8 weeks after the maternal separation, due to the long developmental trajectory of neuron growth in macaques. Here, we added analysis of the adolescent animals and found that in both the 1-WS and 1-MS groups there was a significant reduction in mature neuron cell numbers (MR, 207,353 ± 36,672 versus 1-WS 122,885 ± 10,582, p=0.0110 and 1-MS 118,920 ± 26,670, p=0.0056), a decrease of 40.7% and 42.6% respectively. Immature (DCX+) neurons were also reduced in the maternally separated adolescent groups, but did not reach statistical significance (MR, 530,149 ± 103,977, versus 1-WS, 299,176 ± 19,306, p=0.0563 and 1-MS, 322,540 ± 143,468, p= 0.0635).

**Figure 5:**
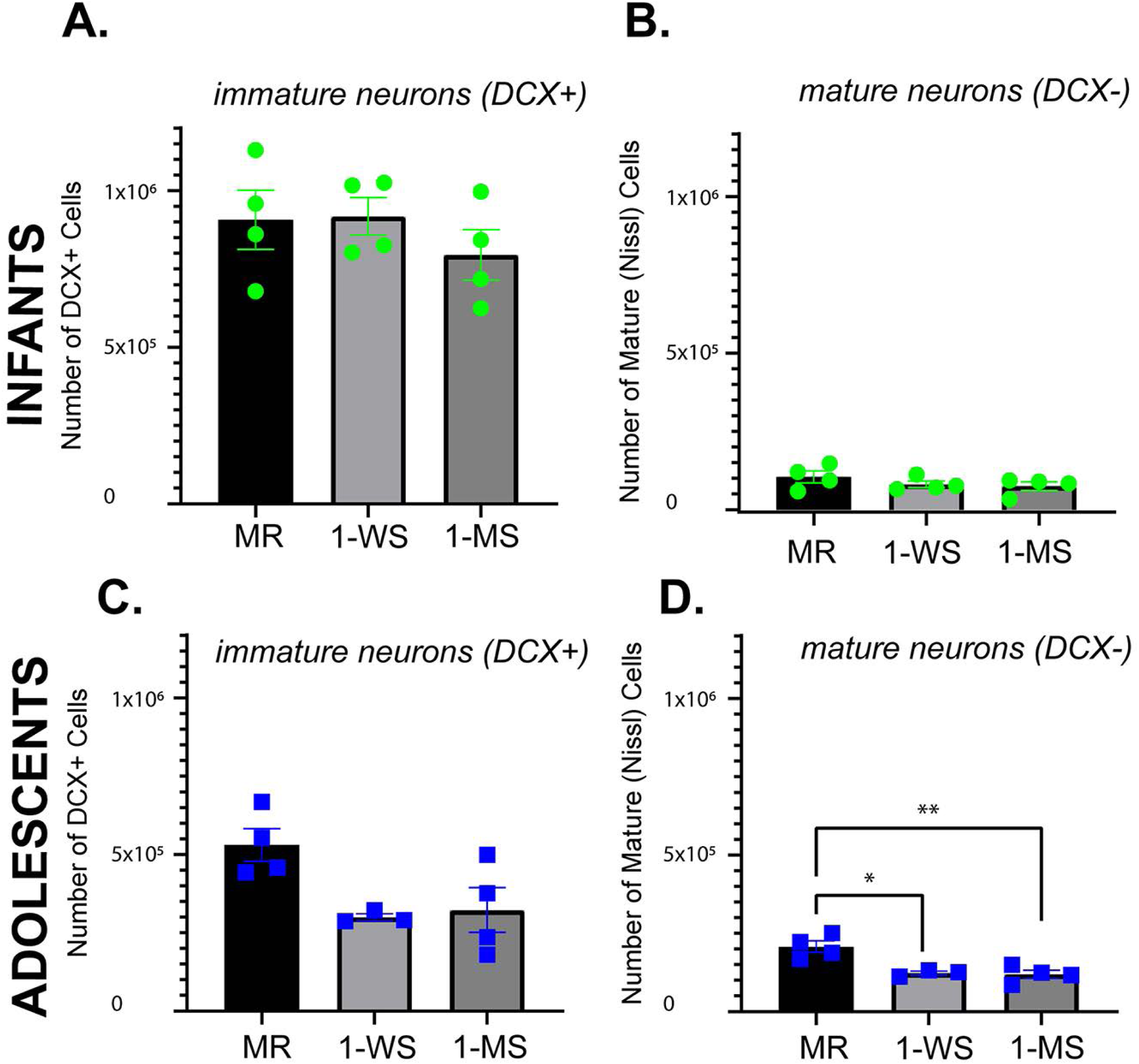
Immature and mature neuron counts in the PL. **A-B.** Infant cohorts (previously published McHale-Matthews et al, shown for comparison, green). By 12 weeks (3-months) of age, there were no significant differences in the number of immature neurons (DCX+) or the number of mature neurons (Nissl) between the MR control, 1-WS, and 1-MS groups **C-D.** Adolescent cohorts (blue dots). By adolescence (4–5 years), the number of immature neurons (DCX+) **(C)** trended lower in the maternally separated groups (1-WS and 1-MS) compared to MR controls but was not significant. Mature neuron (DCX-negative) numbers **(D)** were significantly reduced in 1-WS and 1-MS groups. Statistical analysis was performed using one-way ANOVA with post-hoc testing comparing separation groups to the MR control group within each age stage. Statistical significance: *p < 0.05, **p < 0.01, ***p < 0.001, ****p < 0.0001.

## Discussion

The paralaminar nucleus (PL) is the sole amygdala subregion that retains a lifelong repository of immature neurons which undergo protracted maturation as the animal matures (Avino et al., 2018; Sorrells et al., 2019; Chareyron et al., 2021; McHale-Matthews et al., 2023). In development, maturation of glutamatergic neurons is typically regulated by activity-dependent glutamatergic inputs and neurotrophic factor signaling which stabilize developing synapses (Rakic et al., 1986; Purves, Snider and Voyvodic, 1988; Katz and Shatz, 1996). PL late-developing neurons presumably undergo similar activity-dependent growth, with sustained microglial sculpting of neuronal architecture as neurons develop.

One key finding was that the PL undergoes a pronounced reduction in the number of presynaptic and excitatory post-synaptic elements as well as putative synapses, between infancy and adolescence under normative rearing conditions. These shifts are accompanied by robust increases in microglial volume, phagocytic activity, and synaptic engulfment between infancy and adolescence, consistent with pruning in the adolescent period (Bourgeois, Goldman-Rakic and Rakic, 1994; Petanjek et al., 2011).

A second important finding was that maternal separation disrupts this normative trajectory, due to increased phagocytic activity (microglial volume and CD68 content), premature reductions in synaptic elements, and increased microglial engulfment of synaptic elements in infant animals. Together, these results suggested that accelerated and aberrant microglia-mediated pruning is one mechanism through which early life stress may alter PL cellular maturation, and possibly alter subsequent circuit formation.

A third finding was that while maternal separation profoundly affected synaptic numbers and microglial engulfment in infancy, the consequences for mature neurons emerge later. The DCX+ cells PL have typical features of immaturity including neuritic processes (Fudge, 2004; Chareyron et al., 2021; McHale-Matthews et al., 2023). Increased glutamatergic neuron morphologic maturity in the PL is associated with physiologic maturity (more action potentials, increased axo-somatic synapses)(Alderman et al., 2024). Here, we show maternal separation-related reductions in mature neuron numbers in adolescents, many years after pruning alterations in the infants. While speculative, our results could suggest the possibility that robust changes in synaptic elements induced by maternal separation may have a delayed, but lasting, effect on PL neuron numbers and their development.

### Microglial remodeling and synaptic pruning during normal PL development

The striking reduction in synaptic elements, including contacts, in the PL between infancy and adolescence in typical animals, is consistent with prior reports of synaptic pruning processes described in human and nonhuman primate cortex (Gonzalez-Burgos et al., 2008; Petanjek et al., 2011) and hippocampus (Wei et al., 2012). In normal animals, a large concomitant rise in PL CD68/IBA1 ratios between infancy and adolescence indicated that microglial increases during the interval are not solely attributable to microglial volume expansion but reflect enhanced CD68 enrichment in microglia. Consistent with this, there was an order-of-magnitude increase in engulfed synaptic elements by adolescence, which was detected using two complementary analyses. Together, these data suggest that microglia expand in size and CD68 content during normal development to shape PL circuits.

### Disrupted maternal care induces aberrant pruning primarily in infants

PL microglia undergo a normative developmental shift from amoeboid to more ramified morphologies across infancy and adolescence, consistent with a role in transiently heightened phagocytic activity early in life(Schafer et al., 2012; Schafer, Lehrman and Stevens, 2013). In the PL, we previously found that maternal separation in this same cohort changes this amoeboid to ramified trajectory, resulting in a predominant hyper-ramified state for both infants and adolescents. In this study, we additionally find that maternal separation altered microglia volumes and elevated engulfment of pre- and post-synaptic elements in infancy. These findings support models in which early life stress impacts synaptic modifications microglia actively sculpting neural circuits through synaptic pruning (Tynan et al., 2010; Hinwood et al., 2012; Zhan et al., 2014; Delpech et al., 2016; Reemst et al., 2022) and suggests that early life stress may bias this process toward excessive elimination during infancy. Accelerated microglial-mediated pruning in the infant PL may represent an adaptive response to stress, potentially prioritizing early circuit consolidation or optimizing circuits for survival in adverse contexts (Delpech et al., 2016; Catale et al., 2020).

Notably, stress group differences in both synaptic markers and microglial engulfment were generally absent by adolescence, with one notable exception. Although the individual microglia analysis did not reveal significant changes in PSD95 engulfment, we detected stress-related increases in microglial engulfment of PSD95 elements in adolescence in the ROI neuropil analysis, which mirrored trends in the whole microglia analysis. This may indicate lingering effects of early life stress on excitatory post-synaptic sites (spines), and it is notable that microglia in adolescence had a persistent hyper-ramified appearance(King et al., 2025). Earlier effects on synapse elimination may also have produced downstream effects on spine production and remodeling. In short, even if broader structural measures appear largely ‘normalized’ by adolescence, these changes may contribute to later effects on gene expression and/or connectivity, consistent with extensive evidence linking altered or excessive synaptic pruning during early life to neuropsychiatric disorders (Sekar et al., 2016; Wohleb et al., 2018; Reemst et al., 2022). These findings position the PL as a key site where early life stress and microglial activity converge to confer later vulnerability particularly for amygdala circuits.

### Potential mechanisms connecting early pruning to reduced late-developing neurons

Although we cannot directly connect microglial engulfment in infancy to reduced neuron numbers many years later, the developmental sequence strongly suggests this link. Excessive pruning early in life may limit trophic support needed for neurons to mature, yielding persistent reductions in the adult PL cell population. One mechanism may involve stress-induced stimulation of immature glutamatergic neurons. Stress amplifies glutamatergic pruning responses through purinergic release (Bollinger et al., 2022), and stress-induced neuronal release of colony-stimulating factor 1 (CSF1) similarly promotes phagocytosis (Wohleb et al., 2018). Thus stress-induced neuronal excitation may accelerate microglial pruning mechanisms, leading to a relative deafferentation of immature neurons.

### Disrupted maternal care models: translating between species

Our animal model resulted in similar effects found in many rodent models of maternal separation/disrupted care, such as dendritic spine and synapse loss in the cortex and hippocampus (Review, (Smail and Lenz, 2024). We had similar findings in the infant PL, despite a paradigm tailored for translational studies in primates. As in human societies, monkeys experiencing disrupted maternal care exhibit behavioral and physiological characteristics comparable with those of children experiencing disrupted maternal care including increased display of anxious behaviors, aberrant attachment patterns, changes in adrenal axis regulation and changes in social behavior (Maestripieri et al., 2006; Tottenham et al., 2012; Olsavsky et al., 2013; Howell et al., 2017). When infant monkeys are abandoned in the wild, adolescent and adult females take on caregiving, which involves interacting with and take on care of the infant. Our model is a naturalistic form of disrupted care where the parent is removed, but the rest of the social living situation is retained, as often occurs in humans with death of a parent, illness and poverty. Interestingly, this model induced some of the microglial and synapse changes reported in rodent models of disrupted care.

### Limitations

Out of necessity, our study relied on immunohistochemical markers in fixed tissue, limiting direct assessment of microglial motility, synaptic turnover, or functional interactions. This is a known issue in studying synaptogenesis and microglial engulfment at static timepoint. Moreover, we did not have a ‘curve’ of age groups, and the study was restricted to infant and adolescent timepoints. Nevertheless, we found robust changes in infant PL, using several measures of phagocytosis.

Another potential limitation is related to confocal microscopy. Puncta detected using confocal microscopy in fixed tissue may detect true terminals, but may also detect proteins ‘in transit’ axons or dendrites. Furthermore, ‘contacts’ between presumptive pre- and post-synaptic elements are approximated based on measures from electron microscopic studies. Here, we chose conservative puncta size and ‘contact’ parameters based on data from mature excitatory synapses, since little is known about the developing primate brain. This resulted in high numbers of engulfed SYN1+ and PSD95+ puncta—without synaptic partners--compared to putative contacts. Puncta may reflect elements of nascent (evolving) synapses or non-synaptic proteins ‘in transport’ to the synapse, an issue that is difficult to resolve in fixed tissue (Ahmari, Buchanan and Smith, 2000). Despite these limitations, our data provide a ‘snapshot’ in time, and show robust and consistent shifts in presumptive numbers of synaptic elements, microglial characteristics, and engulfment across age and disrupted early care conditions.

A final limitation was a sample that was predominantly female (19 females, 4 males), precluding analysis of sex-specific effects. Lack of power to detect sex differences limits this study to a ‘first step’, wand will require future studies to add male subjects.

## Conclusion

In summary, maternal separation accelerates synaptic pruning in the infant primate PL, a process mediated by heightened microglial engulfment of synaptic elements. These findings are also associated with a significant decrease in the PL’s population of mature neurons by adolescence, Together, these findings identify the PL as a critical site of vulnerability to early life stress and suggest that microglia-mediated synaptic remodeling may shape subsequent amygdala circuit function.

## Acknowledgements

URMC Center for Advanced Light Microscopy and Nanoscopy (CALMN).

